# High performance genetically-encoded green fluorescent biosensors for intracellular L-lactate

**DOI:** 10.1101/2022.10.19.512892

**Authors:** Giang N. T. Le, Saaya Hario, Kei Takahashi-Yamashiro, Selene Li, Mikhail Drobizhev, Yusuke Nasu, Robert E. Campbell

**Author notes:** Corresponding authors. (Y.N.) and (R.E.C.). These authors contributed equally (Giang N. T. Le, Saaya Hario).

## Abstract

L-Lactate is a monocarboxylate produced during the process of cellular glycolysis and has long been generally considered a waste product. However, studies in recent decades have provided new perspectives on the physiological roles of L-lactate as a major energy substrate and a signaling molecule. To enable further investigations of the physiological roles of L-lactate, we have developed a series of high-performance (Δ*F*/*F* = 15 to 30 *in vitro*), intensiometric, genetically-encoded green fluorescent protein (GFP)-based intracellular L-lactate biosensors with a range of affinities. We evaluated the performance of these biosensors by *in vitro* and live-cell characterization and demonstrated the utility with imaging applications in several cell lines.

## Introduction

Lactate is a monocarboxylate produced from pyruvate by lactate dehydrogenase (LDH) during the process of glycolysis. It can exist as two stereoisomers, L-lactate and D-lactate, with the former being the predominant enantiomer in the human body^1^. L-Lactate has long had a reputation as a metabolic waste product, supported by reports such as a 1929 study that revealed a strong correlation between L-lactate concentration and muscle fatigue in frogs^2^. However, more recent studies have revealed that this unfavorable reputation is undeserved and should be reconsidered^3^. L-Lactate is now known to have many favorable physiological roles as both an energy fuel source^4^ and a signaling molecule^5, 6^, in processes that include memory consolidation^7^, immune response^8, 9^, and neurogenesis^10^. That said, L-lactate is also unfavorably implicated in a number of pathological processes, including inflammation^11, 12^, cancer^13–16^, and neurodegeneration^17^.

Growing evidence that L-lactate has a variety of both favorable and unfavorable physiological roles has inspired efforts to engineer L-lactate-specific genetically encoded fluorescent biosensors. Such biosensors could be powerful tools for studying the concentration dynamics of L-lactate in live cells and tissues. The archetype of such biosensors is the Förster resonance energy transfer (FRET)-based biosensor known as Laconic^18^. Laconic is composed of a cyan fluorescent protein (FP) and a yellow FP fused to the termini of the L-lactate-binding transcription factor LldR. It has been effectively used in a variety of applications including investigations of the Warburg effect^18, 19^ and the astrocyte–neuron L-lactate shuttle (ANLS) hypothesis^20, 21^. Drawbacks of Laconic include its very small ratiometric response and its use of two colors of FP which limits the opportunities for multiplexed imaging with other colors of biosensor.

Relative to FRET-based indicators, single FP-based indicators can typically be engineered to have much larger intensiometric fluorescent responses and are suitable for multiplexed imaging applications. In efforts aimed at realizing these advantages, a number of single FP-based L-lactate biosensors have been engineered in recent years^22–27^. These single FP-based biosensors share a general design in which the L-lactate-binding protein is genetically linked to an FP such that the binding protein is located close to the chromophore^28^. Binding to L-lactate causes a conformational change that changes the chromophore environment and, consequently, the intensity of the fluorescence. Representative examples include the eLACCO1.1 and R-eLACCO2 biosensors for extracellular L-lactate, both based on the TTHA0766 L-lactate binding protein^26, 27^. These biosensors offer high performance in the extracellular milieu but since they have a strict requirement for Ca^2+^ (10s of μM), they do not function in the cytosolic environment where the Ca^2+^ concentration is < 1 μM. The existing LldR-based biosensors do not suffer from this Ca^2+^-dependence, but unfortunately have relatively limited fluorescence responses towards L-lactate^22, 23^.

Here we report a series of three new single GFP-based biosensors for L-lactate that overcome the limitations of previously reported biosensors and have affinities that span the physiological concentration range of L-lactate. These biosensors, designated iLACCO1, iLACCO1.1, and iLACCO1.2, are the final product of extensive directed evolution and structure-guided mutagenesis. As we demonstrate in this work, the iLACCO biosensors provide outstanding performance that greatly facilites imaging of intracellular L-lactate dynamics in mammalian cells.

## Results

### Development of a genetically encoded L-lactate biosensor, iLACCO1

An initial prototype of the L-lactate biosensor was constructed by inserting circularly permuted green fluorescent protein (cpGFP), derived from iGluSnFR^29^, into the L-lactate binding domain (LBD) of the *Escherichia coli* LldR transcriptional regulator protein^30^ at various solvent accessible positions (Fig. 1a). Three residue linkers, DWS at the N-terminus (first linker) and NDG at the C-terminus (second linker) of cpGFP, were introduced to connect the cpGFP domain to the LldR-LBD (Fig. 1b). These linkers are connected to the two “gate post” residues of cpGFP (His145 and Phe148 as numbered in GFP; His110 and Phe352 as numbered in Supplementary Fig. 1). These gate post residues have been proposed to play an important role in cpGFP-based biosensors^28^. Insertion sites in LldR-LBD were chosen manually based on an AlphaFold model of the structure^31, 32^, and were solvent-exposed sites that were considered likely to undergo L-lactate-dependent conformational changes. A total of 11 different insertion sites, located in three different solvent accessible loops, were tested.

**Fig. 1.**
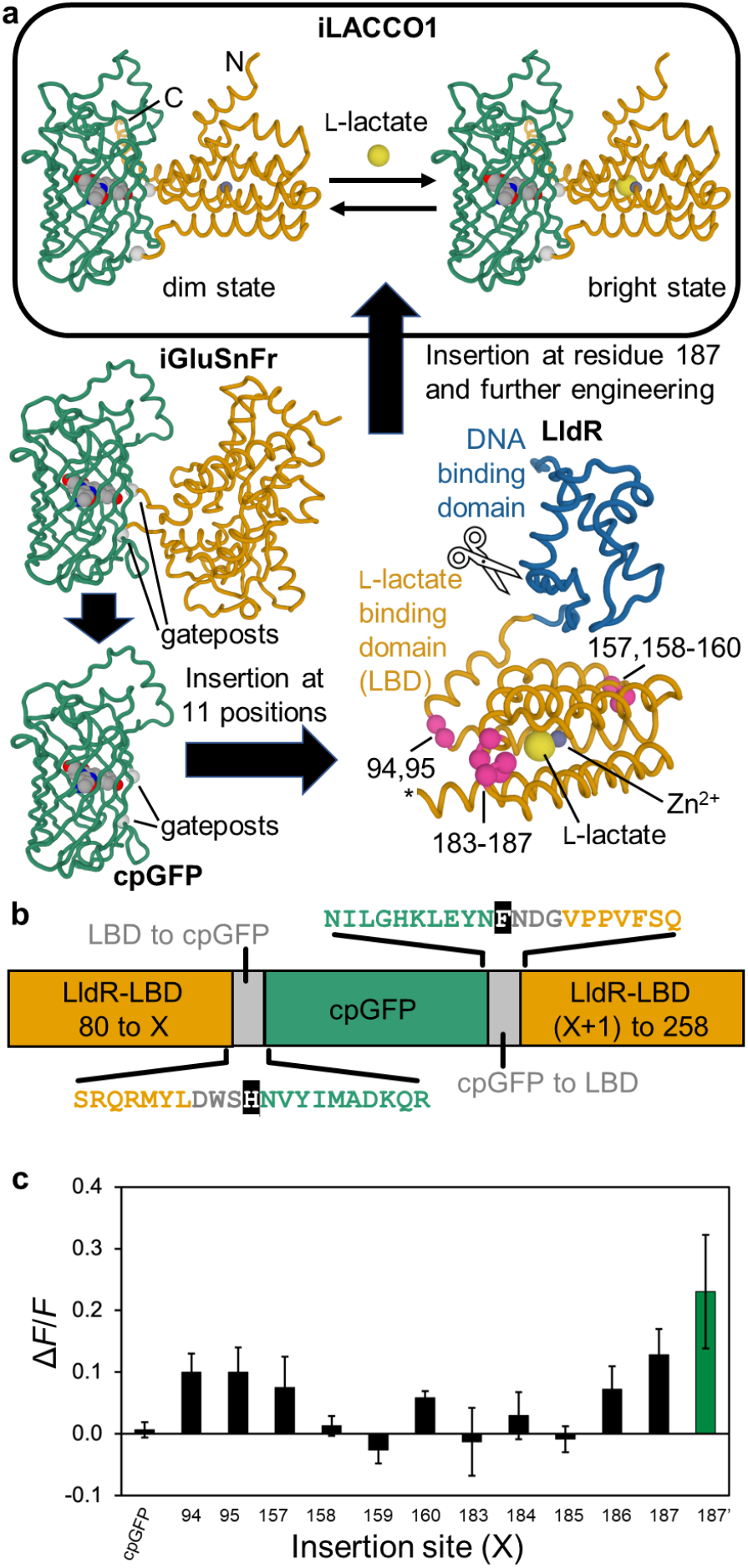
iLACCO design strategy. (**a**) Schematic representation of the overall strategy used to engineer iLACCO1. Structures shown are AlphaFold^31, 32^ models of iLACCO1, iGluSnFr^29^, cpGFP, and *E. coli* LldR. The Zn^2+^ (purple sphere) and L-lactate (yellow sphere) were positioned based on a superposition with the sialic acid-binding homolog NanR (PDB ID: 6ON4)^44^. Gate post residues demarcate the beginning and end of the cpGFP domain. Pink spheres represent insertion sites that were initially tested. To remove the N-terminal DNA-binding domain, the region of DNA encoding the first 79 residues of LldR was removed. (**b**) Schematic representation of the 11 insertion site variants initially tested. Linker regions are represented in gray. Gate posts are represented with white text on a black background. (**c**) Δ*F*/*F* of each prototype biosensor, where cpGFP is inserted at the site of LldR-LBD indicated on the horizontal axis. A variant with insertion of cpGFP at site 187, which also had a point mutation in the second linker (NDG to NEG) (187’; green bar) gave the largest absolute value of Δ*F*/*F* and so this protein was designated iLACCO0.1. *n* = 3 technical replicates, mean ± s.d.

In the process of cloning a variant with cpGFP inserted at position 187, a single colony (designated 187’) was found to be much brighter than others. This clone was later determined to have a point mutation in the second linker (NDG to NEG). This variant had the largest change in fluorescence intensity (Δ*F*/*F* = (*F*_max_ - *F*_min_) / *F*_min_) upon adding L-lactate, and exhibited a direct response (fluorescence increase upon binding) to L-lactate of Δ*F*/*F* = 0.23 (Fig. 1c). This variant was designated iLACCO0.1 and used as the template for further engineering.

To further develop iLACCO0.1 to obtain variants with larger absolute Δ*F*/*F*, we first optimized the linker lengths (Supplementary Fig. 2). Site-directed mutagenesis was used to obtain variants with partial or total deletions of both the first and second linkers (Supplementary Fig. 2b). The variant with all three amino acids of the first linker deleted, and all residues of the second linker retained, was determined to have the greatest response to L-lactate and was designated iLACCO0.2. This variant exhibited an inverse response (fluorescence decrease upon binding) with Δ*F*/*F* = −0.55 (Supplementary Fig. 2c).

To further improve the fluorescence response, we next optimized the linker sequences and then performed directed evolution of the whole gene. Since the first linker had been deleted, three amino acids (Met107, Tyr108, and Leu109) of LldR-LBD, adjacent to the gate post residue His110, were considered the new N-terminal linker. We constructed and screened a series of libraries in which pairs of residues of the N-terminal and C-terminal linkers were sequentially randomized. Successive screening of these libraries led to the discovery of iLACCO0.4 with a direct response (Δ*F*/*F* = 1.3) to L-lactate. Further optimization was performed using directed evolution as shown schematically in Fig. 2a. Eleven rounds of directed evolution led to the final iLACCO1 variant with Δ*F*/*F* of 20 under the screening conditions (*i.e*., crude protein extract in B-PER solution) (Fig. 2b). Relative to iLACCO0.1, iLACCO1 contains 21 point mutations (Fig. 2c,d and Supplementary Fig. 1).

**Fig. 2.**
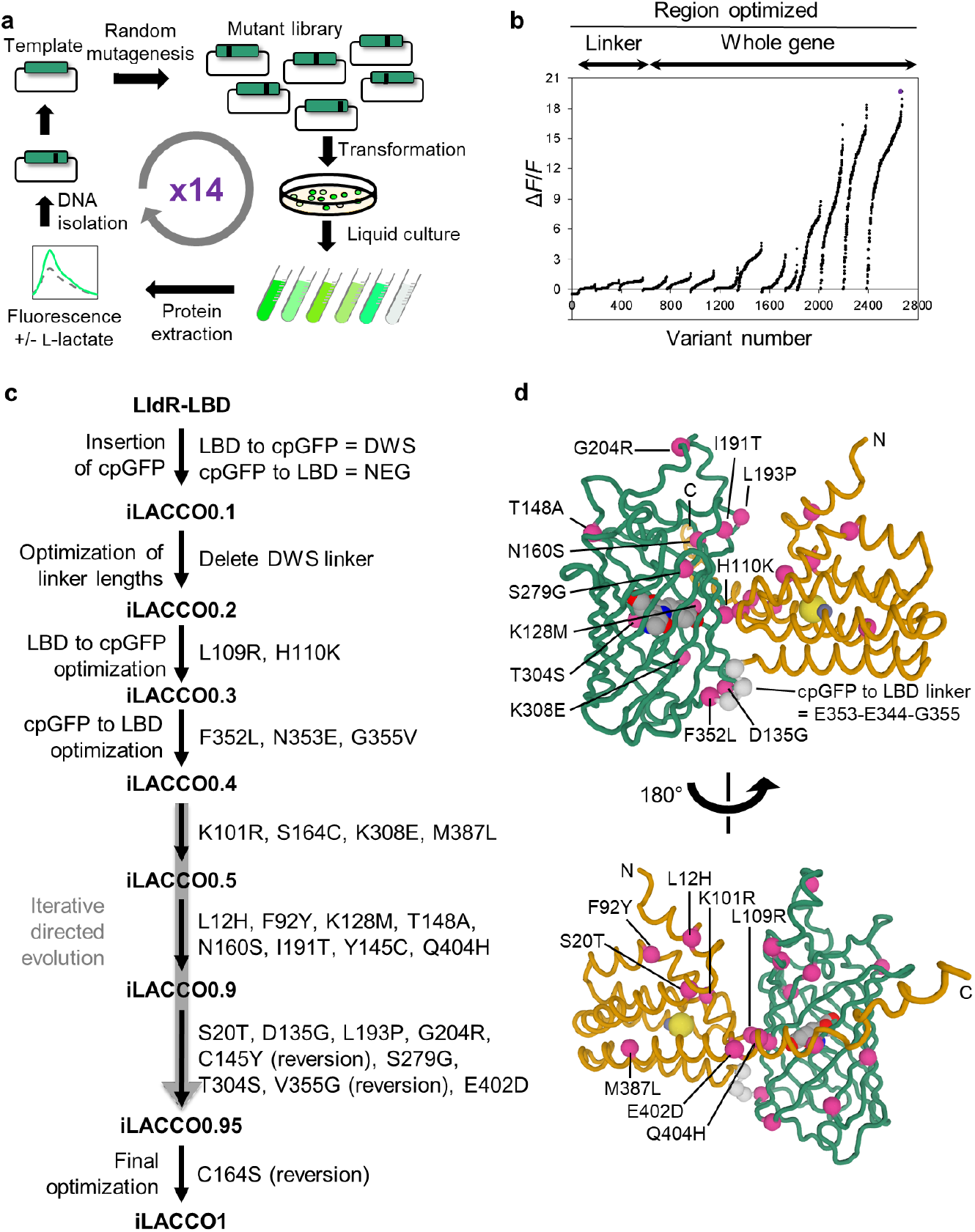
Directed evolution of iLACCO1. (**a**) Schematic of directed evolution workflow. Starting from the template of iLACCO0.4, the full-length gene was randomly mutated by error-prone PCR and the resulting library was used to transform *E. coli*. Bright colonies were picked and cultured, and Δ*F*/*F* upon addition of 10 mM L-lactate was determined using crude protein extracts. The genes encoding the variants with the highest Δ*F*/*F* were used as the template for the next round. (**b**) Δ*F*/*F* rank plot representing all proteins tested during the directed evolution. For each round, tested variants are ranked from lowest to highest Δ*F*/*F* value from left to right. (**c**) Lineage of iLACCO variants from LldR-LBD. (**d**) Modeled structure^31, 32^ of iLACCO1 with the position of mutations indicated.

### *In vitro* characterization of iLACCO1

Characterization of key photophysical and biochemical properties of purified iLACCO1 revealed it to be a high-performance biosensor. In the presence of L-lactate, iLACCO1 exhibits absorbance peaks at 400 nm and 493 nm (Fig. 3a), corresponding to the neutral (protonated) and the anionic (deprotonated) forms of chromophore, respectively. Kinetic measurements for iLACCO1 treated with L-lactate revealed a rapid increase in absorbance (*t*_0.5_ = 0.42 s) to approximately 60% of its maximum value, followed by a much slower increase (*t*_0.5_ = 16.4 minutes) (Supplementary Fig. 3). The excitation spectrum of iLACCO1 in the presence of L-lactate has a maximum at 493 nm, consistent with the absorbance spectrum, and the emission maximum is 510 nm (Fig. 3b). Purified iLACCO1 has a Δ*F/F* of 30 (Fig. 3b) and an apparent dissociation constant (*K*_d_) of 361 µM (Fig. 3c) for L-lactate at pH 7.2. iLACCO1 also exhibits p*K*_a_ values of 7.4 and 8.8 in the presence and absence of L-lactate, respectively (Fig. 3d). The two-photon spectrum of iLACCO1 reveals that the excitation maximum of the L-lactate bound state is 928 nm (where the 2P absorption is dominated by the anionic form) with brightness of *F*_2_ = 7.3 GM, 1GM = 10^-50^ cm^4^s (*F*_2_ ≃ σ_2,A_ × φ_A_ × ρ_A_, where the two-photon absorption cross section, σ_2,A_, the fluorescence quantum yield, φ_A_, and the relative fraction ρ_A_, - all correspond to the anionic form of chromophore). The Δ*F*_2_*/F*_2_ value ranges from 13.0–14.7 in the 928–1000 nm wavelength range (Fig. 3e).

**Fig. 3.**
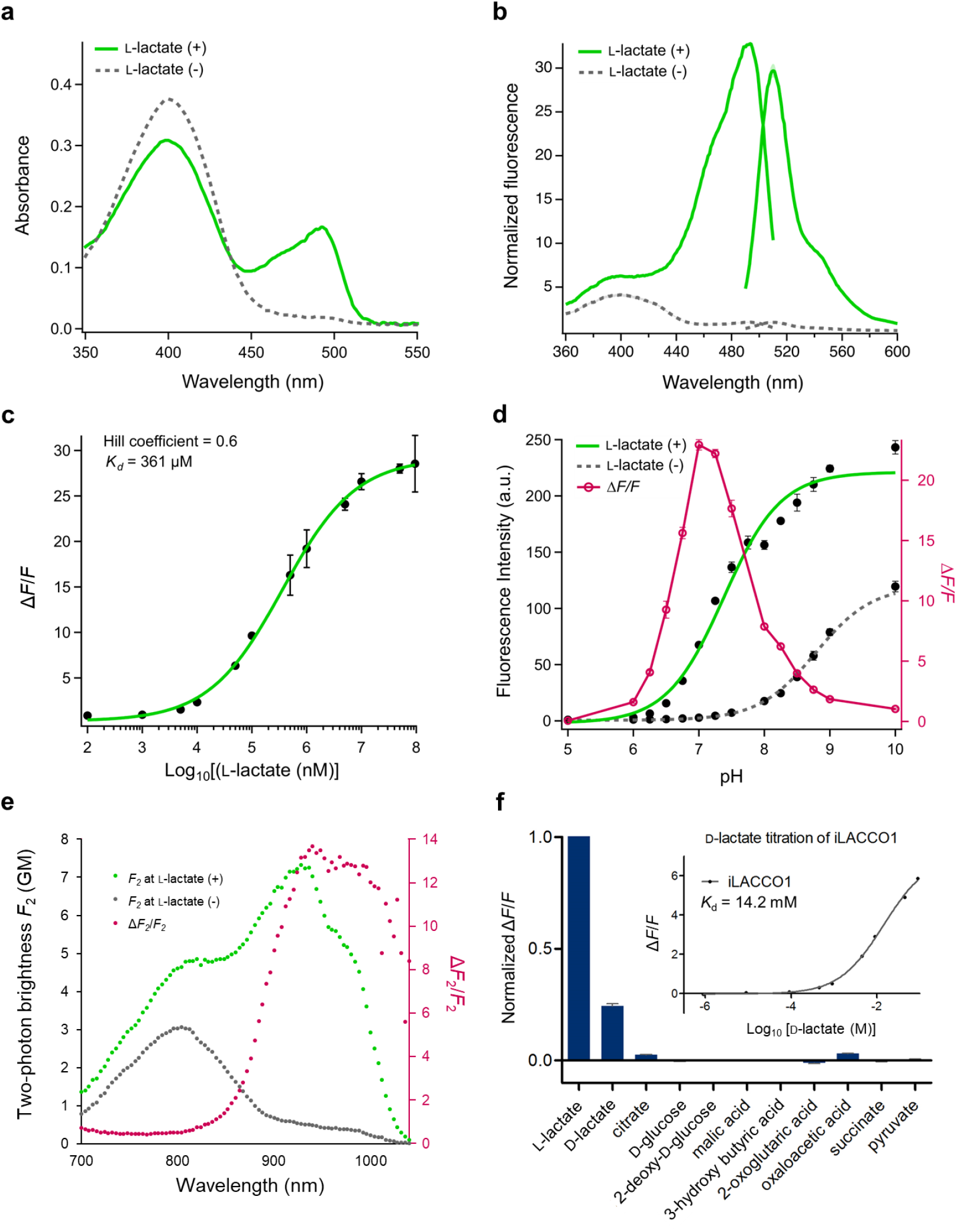
*In vitro* characterization of iLACCO1. (**a**) Absorbance spectra of iLACCO1 in the presence (10 mM) and absence of L-lactate. (**b**) Excitation (emission at 570 nm) and emission spectra (excitation at 450 nm) of iLACCO1 in the presence (95 mM) and the absence of L-lactate. (**c**) Dose-response curve of iLACCO1 for L-lactate. *n* = 3 technical replicates (mean ± s.d.). (**d**) pH titration curve of iLACCO1 in the presence (10 mM) and the absence of L-lactate. *n* = 3 technical replicates (mean ± s.d.). (**e**) Two-photon excitation spectra of iLACCO1 in the presence (10 mM) (represented in green dots) and absence of L-lactate (represented in gray dots) shown with the GM values label at left Y-axis. Δ*F*_2_/*F*_2_ is the ratio of the two-photon excitation spectra (represented in magenta dots) labeled at right Y-axis. (**f**) Molecular specificity of iLACCO1 and dose-response curve of iLACCO1 for D-lactate. *n* = 3 technical replicates (mean ± s.d.).

*In vitro* testing revealed that iLACCO1 is highly specific for L-lactate and has negligible fluorescence response to the structurally similar molecules and representative metabolites listed in Fig. 3f. iLACCO1 does exhibit a substantial fluorescence response to its enantiomer, D-lactate, though with a 40× lower affinity (*K*_d_ = 14.2 mM; Fig. 3f **inset**). Notably, the physiological concentration of D-lactate in plasma is 10s of µM^33^, which is hundreds of times lower than the apparent *K*_d_ of iLACCO1 for D-lactate. Note that the concentration of D-lactate was mistakenly stated to be in the nM range in an oft-cited 2005 review^34^. Due to its lower affinity and the lower physiological concentration, D-lactate is unlikely to interfere with iLACCO-based measurements of L-lactate concentration.

### Development of iLACCO variants with different affinities

Range of physiological concentration of intracellular L-lactate depends on cell types^20^. In parallel with the process of directed evolution which ultimately produced iLACCO1, we also undertook the development of variants with different affinities for their wider applicability. Based on the X-ray crystal structure of LldR from *Corynebacterium glutamicum*^30^, we constructed a homology model of LldR from *Escherichia coli* using the M4T Server ver. 3.0 (ref. 35). Based on the homology model, the L-lactate-binding cavity of LldR is lined with hydrophobic residues that likely interact with L-lactate, and charged residues that likely interact with a zinc ion that coordinates with L-lactate^30^. Focusing on the hydrophobic interactions, we designed a series of ten conservative mutations (E39Q, D69E, M89Q, F93Y, L96I, V100T, V100I, L364I, V393T, and V393) that could potentially have an effect on L-lactate binding affinity (Supplementary Fig. 4a). Each of these ten mutations was individually introduced by site-directed mutagenesis into iLACCO0.5, and affinities for L-lactate were measured. Mutations which changed the affinity but did not abolish the fluorescence response (V100T, V100I, V393T, and V393I) were selected for further investigation (Supplementary Fig. 4b). Testing these mutations in the context of iLACCO0.9 enabled us to identify Val393Ile as the best mutation for decreased affinity (*K*_d_ = 4.55 mM), and Val100Ile as the best mutation for increased affinity (*K*_d_ = 16.9 µM) (Supplementary Fig. 4c). Finally, these mutations were introduced into iLACCO1 to produce iLACCO1.1 (iLACCO1 V393I; low affinity) and iLACCO1.2 (iLACCO1 V100I; high affinity) (Fig. 4a,b).

**Fig. 4.**
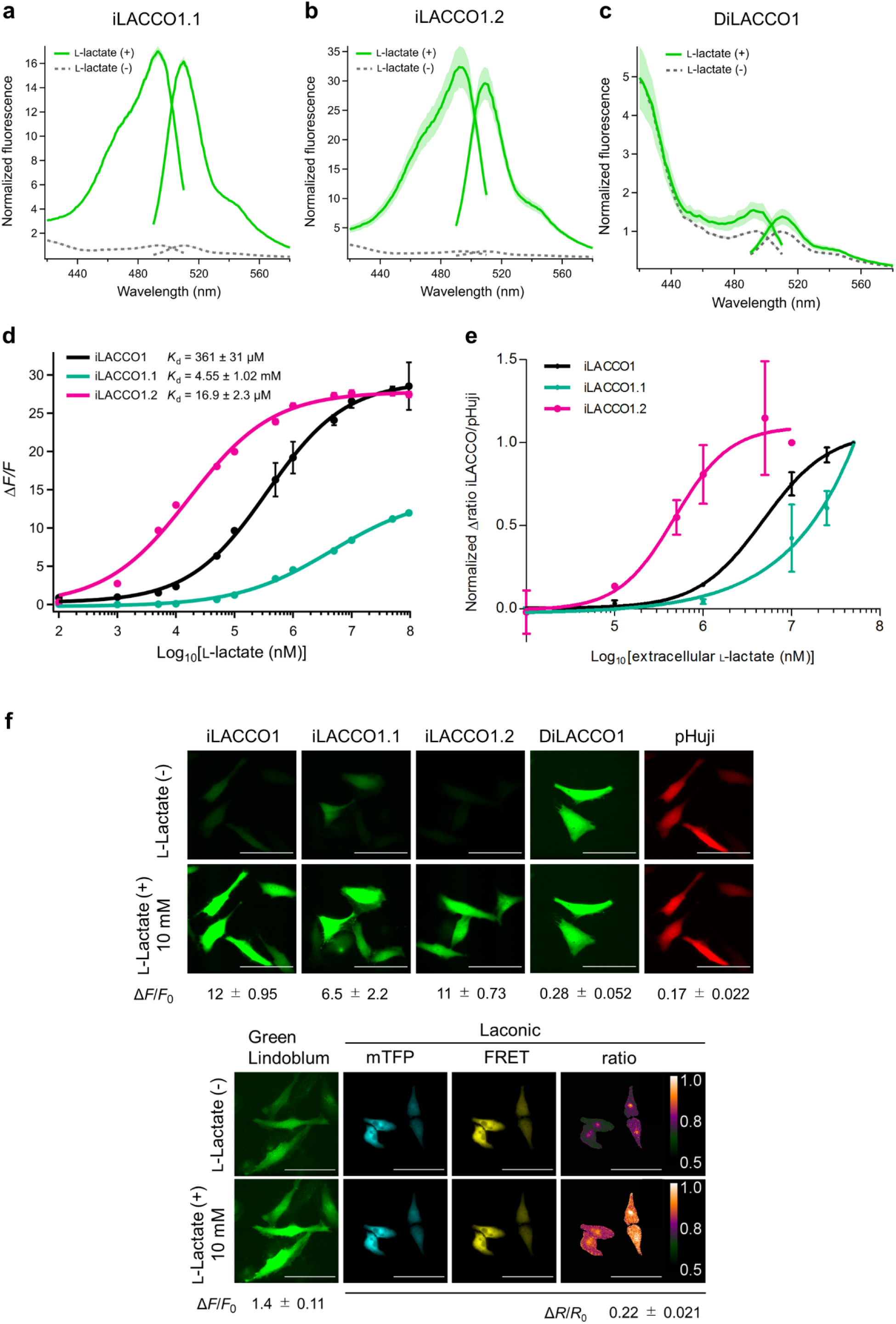
Affinity tuning of iLACCO1. (**a**-**c**) Excitation and emission spectra of iLACCO variants in the presence (95 mM) and the absence of L-lactate. *n* = 3 technical replicates (mean ± s.d.). (**d**) Dose-response curves of purified iLACCO1 variants upon treatment with L-lactate. *n* = 3 technical replicates (mean ± s.d.). (**e**) Dose-response curves of HeLa cells expressing iLACCO1 variants in response to treatments with extracellular L-lactate, as measured using flow cytometry. iLACCO1, 1.1, and 1.2 gave 50% of their maximal response at treatment concentrations of 4.8 mM, > 10 mM, and 0.47 mM, respectively. *n* = 3 from independent experiments (mean ± s.d.), and around 1.0 × 10^5^ cells were analyzed for each independent experiment. (**f**) Fluorescence images of HeLa cells expressing iLACCO variants, pHuji^36^, Green Lindoblum^22^, and Laconic^18^ in the presence (10 mM) and absence of L-lactate. Δ*F*/*F*_0_ and Δ*R*/*R*_0_ are calculated from cells in the images shown. Scale bars represent 100 µm. *F*_0_ and *R*_0_ are determined as average fluorescence intensity of 15 data points before addition of any reagent.

As with most genetically encoded biosensors, the fluorescence of iLACCO1 and its variants is highly sensitive to pH changes within the physiological range. Control biosensors that do not respond to the target of interest, but do exhibit a similar pH sensitivity as the biosensor (in one state or the other), are useful tools for helping researchers distinguish a true response from a pH-induced artifact. Having found that mutations of Val100 and Val393 can affect L-lactate binding, further mutations were introduced in these sites. The combination of Val100Ala and Val393Ala abolished the fluorescence response to L-lactate. This variant, designated ‘<u>d</u>ead’ iLACCO1 (DiLACCO1), showed pH dependence that was similar to the lactate-free state of iLACCO variants and was used as the control biosensor (Figs. 3d, 4c and Supplementary Fig. 5). The dose-response curves of the iLACCO variants towards L-lactate are shown in Fig. 4d, and the two-photon spectra of iLACCO1.1 and iLACCO1.2 are shown in Supplementary Fig. 6. The photophysical and biochemical properties are summarized in Supplementary Table 1.

### Characterization of iLACCO1 variants in mammalian cells

iLACCO1 variants were characterized in HeLa cells using flow cytometry and fluorescence microscopy. We cloned each of the iLACCO genes into a mammalian expression vector with a CMV promoter. As a spectrally orthogonal control for possible pH changes, to the 3’ end of the iLACCO gene we appended the gene for the red fluorescent protein (RFP)-based pH biosensor pHuji^36^, separated by a self-cleaving P2A sequence^37^. To assess the functions of iLACCO1 variants based on large populations of cells, flow cytometry was conducted with cells that were expressing iLACCO variants and had been treated with various concentrations of L-lactate. Cells were treated with iodoacetic acid to stop intracellular L-lactate production, nigericin to clamp the pH, and rotenone to block mitochondrial metabolism. These conditions have previously been employed for characterization of an L-lactate biosensor^18^. Based on the dose-response curves of iLACCO-expressing cells, iLACCO1, 1.1, and 1.2 gave 50% of their maximal response at treatment concentrations of 4.8 mM, > 10 mM, and 0.47 mM L-lactate, respectively (Fig. 4e).

As another validation of the responses of the iLACCO variants in cells, we performed fluorescence microscopy of HeLa cells that were treated identically to the cells in the cytometry experiments. For the cells shown in Figure 4f, the observed Δ*F*/*F*_0_ values were 12 ± 0.95 for iLACCO1, 6.5 ± 2.2 for iLACCO1.1, 11 ± 0.73 for iLACCO1.2, 0.28 ± 0.052 for DiLACCO1, and 1.4 ± 0.11 for Green Lindoblum. For Laconic, the Δ*R*/*R*_0_ was 0.22 ± 0.021 and for the pHuji pH indicator, the Δ*F*/*F*_0_ was 0.17 ± 0.022. Based on this data, we conclude that the iLACCO series of biosensors retains high performance in cells.

### Imaging applications of iLACCO variants in mammalian cells

To investigate the utility of iLACCO variants for monitoring of L-lactate concentration in mammalian cells, we carried out imaging experiments using HeLa and HEK293 cells in several conditions. As shown in Fig. 5a, cells were cultured in Dulbecco’s modified Eagle medium (DMEM) with high glucose (25 mM) and then 2–3 hours before imaging the media was switched to DMEM with no glucose. During imaging, glucose was added to the media (*t* = 0) to a final concentration of 5 mM. We expected this treatment to cause an increase in intracellular L-lactate concentration and a corresponding increase in iLACCO fluorescence intensity. In both cell lines, the high affinity iLACCO1.2 gave the largest increase in fluorescence. iLACCO1 gave a substantially smaller change and both iLACCO1.1 and DiLACCO1 gave negligible changes (Fig. 5b-e). Assuming that the *K*_d_ values measured with purified protein (Fig. 4d) are retained in the intracellular environment, these results are consistent with a baseline L-lactate concentration of 1 μM or less under starvation conditions, increasing to 10-100 μM upon treatment with 5 mM glucose.

**Fig. 5.**
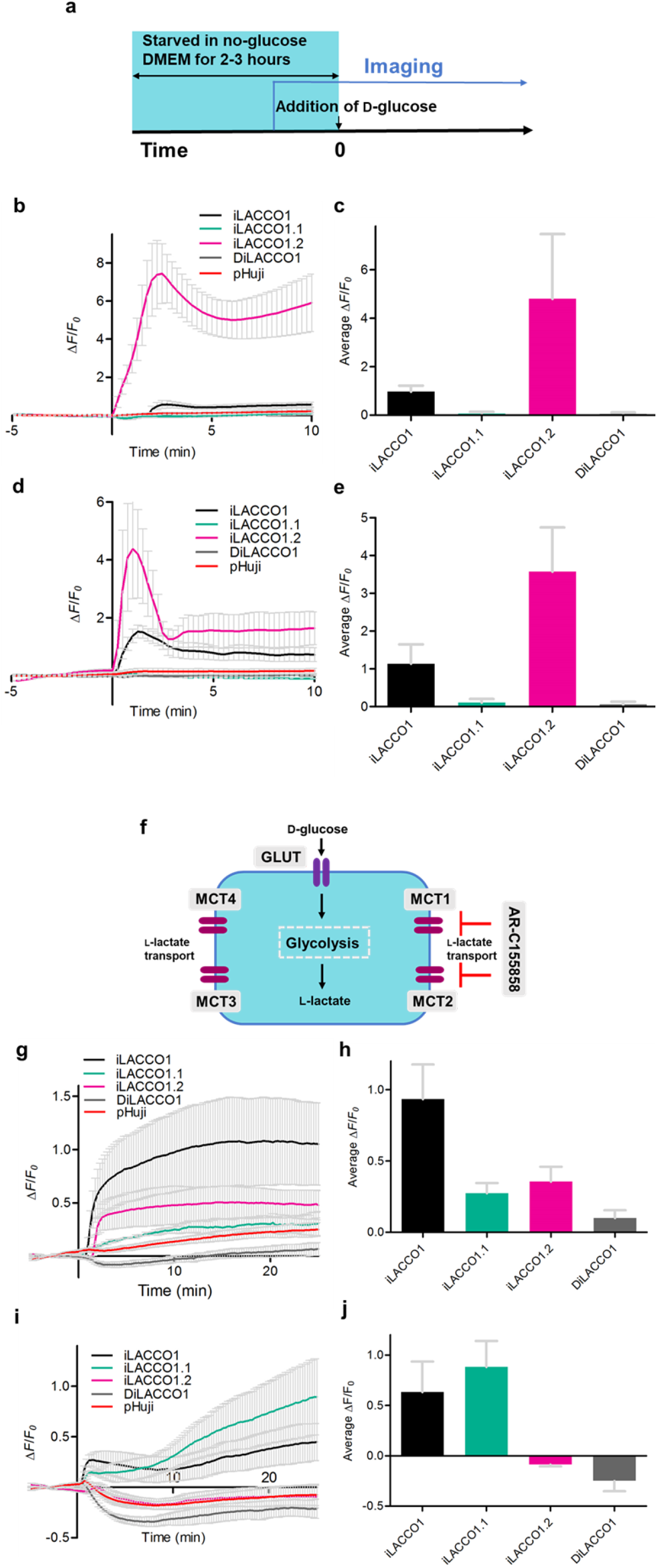
Characterizations of iLACCO variants in live mammalian cells. (**a**) Schematic of imaging condition of starvation experiment. HeLa and HEK293 cells were starved in no-glucose medium for 2–3 hours and were treated with final concentration of 5 mM of D-glucose at time = 0. Glucose-induced changes of intracellular L-lactate concentration were observed with iLACCO1 variants expressed in HeLa (data shown in **b** and **c**) and HEK293 (data shown in **d** and **e**) cells. (**b, d**) Representative time courses show mean ± s.d. iLACCO1 (black, *n* = 6 and 3 cells for HeLa and HEK293, respectively), iLACCO1.1 (green, *n* = 5 and 2 cells for HeLa and HEK293, respectively), iLACCO1.2 (pink, *n* = 5 and 3 cells for HeLa and HEK293, respectively), DiLACCO1 (gray, *n* = 5 and 4 cells for HeLa and HEK293, respectively) from single independent experiment and pHuji (red, *n* = 21 and 12 cells for HeLa and HEK293, respectively). (**c, e**) Bar graphs show the mean ± s.d of maximum Δ*F*/*F*₀ determined as the peak values within 5 minutes after addition of D-glucose. iLACCO1 = (23, 6), iLACCO1,1 = (21, 6), iLACCO1.2 (20, 5), and DiLACCO1 = (10, 3), where (x, y) = (number of cells in total, number of independent experiments). (**f**) Schematic of MCT1,2 inhibitor experiment. Fluorescence intensity change was observed during the addition of AR-C15585 in HeLa (data shown in **g** and **h**) and HEK293 (data shown in **i** and **j**) cells. (**g, i**) Fluorescence response of iLACCO1 variants expressing HeLa and HEK293 cells upon treatment of MCT1,2 inhibitor AR-C155858. AR-C155858 (final 1 μM) was added at 0 min under high glucose (25 mM) condition. Mean ± s.d. iLACCO1 (black, *n* = 4 and 4 cells for HeLa and HEK293, respectively), iLACCO1.1 (green, *n* = 2 and 5 cells for HeLa and HEK293, respectively), iLACCO1.2 (pink, *n* = 3 and 6 cells for HeLa and HEK293, respectively), DiLACCO1 (gray, *n* = 4 and 4 cells for HeLa and HEK293, respectively) from single independent experiment, and pHuji (red, *n* = 13 and 20 cells for HeLa and HEK293, respectively). (**h, j**) Bar graphs show the mean ± s.d values of Δ*F*/*F*_0_ during time = 15 to 25 mins. iLACCO1 = (22, 5), iLACCO1,1 = (16, 5), iLACCO1.2 (17, 4), and DiLACCO1 = (15, 5), where (x, y) = (number of cells in total, number of independent experiments).

To further investigate the utility of iLACCO variants for monitoring of L-lactate concentration in mammalian cells, we examined the effect of an inhibitor of L-lactate flux through membrane transporters. AR-C155858 is a specific inhibitor^38^ of proton-coupled monocarboxylate transporters 1 and 2 (MCT1, MCT2), which transport L-lactate plus a proton across membranes^39^ (Fig. 5f). We treated HeLa cells and HEK293 cells, transfected with iLACCO variants and cultured with high glucose, with this inhibitor and compared their fluorescence intensity changes. In HeLa cells, iLACCO1 (Δ*F*/*F*_0_ ∼ 0.89) showed a substantial increase in fluorescence intensity, while iLACCO1.1 (Δ*F*/*F*_0_ ∼ 0.25) and iLACCO1.2 (Δ*F*/*F*_0_ ∼ 0.28) had smaller increases by time = 15 to 25 mins (Fig. 5g, h). Assuming the *K*_d_ values measured with purified protein (Fig. 4d), these results are consistent with an initial concentration of L-lactate in the range of 100s of μM, increasing to 1 mM or greater upon treatment with AR-C155858. Both iLACCO1 and the higher affinity iLACCO1.2 variant were observed to respond substantially faster than the lower affinity iLACCO1.1 variant. In HEK293 cells, iLACCO1.1 exhibited the largest increase in fluorescence (Δ*F*/*F*_0_ ∼ 0.87), with iLACCO1 exhibiting a smaller but still substantial increase (Δ*F*/*F*_0_ ∼ 0.64). In contrast, iLACCO1.2 did not exhibit any substantial change (Fig. 5i, j). These results are consistent with HEK293 cells having a higher baseline concentration (>1 mM) of intracellular L-lactate than HeLa cells, under high glucose conditions.

## Discussion

We have developed a series of intensiometric, genetically-encoded, high performance, single GFP-based biosensors for intracellular L-lactate, which we have designated as the iLACCO series. The initial prototype of this series was obtained by the insertion of cpGFP into a loop of the L-lactate binding domain (LBD) of the *Escherichia coli* LldR transcriptional regulator protein. Starting from this prototype we undertook extensive linker optimization and directed evolution to arrive at iLACCO1 with a Δ*F*/*F* of ∼30 upon binding to L-lactate. Notably, this is one of the highest fluorescence responses ever achieved for a single GFP-based biosensor for a ligand other than Ca^2+^ (ref. 28). Other recently reported GFP-based L-lactate biosensors have Δ*F*/*F* values of <1, 3.0, 4.2, and 6.0 (refs. 22–24,26).

Site-directed mutagenesis of residues lining the modeled L-lactate binding pocket of the LldR domain resulted in variants with lower (iLACCO1.1) and higher (iLACCO1.2) affinity and high Δ*F*/*F* values of 15 and 28, respectively. Between these three variants, detection of L-lactate concentrations in the range of ∼100 nM to ∼100 mM should be feasible in principle. The imaging applications with HeLa cells and HEK293 cells clearly demonstrates the utility of a series of iLACCO with different affinities that covers a wide dynamic range. Based simply on which affinity variant(s) gave the largest responses in a particular imaging experiment, we could make robust order-of-magnitude estimates of the L-lactate concentration changes, assuming the the *K*_d_ values of in cells are the same as for purified proteins.

For single GFP-based biosensors, the fluorescence signal originating from the GFP is modulated by changes in the chromophore environment that occur as a result of conformational changes associated with ligand binding^28^. Commonly, these changes in the chromophore environment cause a shift in the p*K*_a_ of the chromophore, leading to a change in the relative populations of the dim protonated state (phenol) and the bright deprotonated state (phenolate). The iLACCO series appears to employ just such a response mechanism. The absorbance spectrum of iLACCO1 in the absence of L-lactate reveals that the chromophore exists mostly in the protonated state (absorption peak at 400 nm) at neutral pH. Upon addition of L-lactate, the chromophore partly converts to the deprotonated form (absorption peak at 493 nm) (Fig. 3a). This change is consistent with the observed shift in the p*K*_a_ of the iLACCO1 chromophore from 8.78 for the unbound state to 7.38 for the bound state (Fig. 3d). This increase in the deprotonated state upon binding to L-lactate is the major contributing factor to the response of the biosensor. Notably, a large fraction of the protein remains in the protonated state even in the presence of L-lactate, suggesting that there is room for achieving much larger fluorescence responses, and higher brightness of the bound state, with further engineering.

Although an experimental atomic structure of iLACCO1 is not available, we can refer to an Alphafold model^31, 32^ of the protein and speculate on the molecular interactions that may be responsible for the L-lactate-dependent p*K*_a_ shift. Generally speaking, a shift from a higher to a lower p*K*_a_ could be attributable to a gain of new interactions that stabilize the deprotonated phenolate state (e.g., interaction with a positively charged group), or a loss of interactions that stabilize the protonated phenol state (e.g., interaction with a negatively charged or hydrophobic group). Intriguingly, the N-terminal gate post of iLACCO1 and its preceding residue are both positively charged (His110Lys and Leu109Arg, respectively). Furthermore the C-terminal gate post, and its following residue are both negatively charged (Asn353Glu and Glu354, respectively). Accordingly, we tentatively suggest that the L-lactate dependent conformational change in the LBD is being propagated through the linkers to the cpGFP domain, leading to new interactions with the side chains of residues 109–110, and/or the loss of interactions with side chains of residues 353–354 (Supplementary Fig. 7). Further studies using X-ray crystallography and molecular dynamics simulations will likely be necessary to gain a better understanding of the fluorescence response mechanism.

A common disadvantage of most single GFP-based biosensors is pH sensitivity and the iLACCO series is not an exception. The p*K*_a_ values for the L-lactate bound states of iLACCO1, iLACCO1.1, and iLACCO1.2 are 7.38, 7.65, and 6.76 respectively, values that are all in the physiological pH range. This pH sensitivity is potentially and particularly problematic for L-lactate biosensors, because the MCTs transport L-lactate plus a proton and so L-lactate flux is necessarily associated with changes in pH. Specifically, L-lactate influx should be associated with a decrease in cytosolic pH, and L-lactate efflux should be associated with an increase in cytosolic pH. Fortunately, with respect to iLACCO fluorescence intensity, the effect of decreasing pH (decreased fluorescence intensity) is opposite to the effect of increased L-lactate (increased fluorescence intensity). For this reason, decreases in pH should not generally confound the qualitative interpretation of results obtained from iLACCO imaging, though increases in pH could. For this reason, we recommend that coexpression of a spectrally distinct pH biosensor, such as pHuji, should be routinely done when using the iLACCO series. Decreased sensitivity to pH changes could be an important feature to engineer into future iLACCO variants. As precedent for such engineering, the L-lactate biosensor designated LiLac^25^ has been optimized for fluorescence lifetime imaging and minimal pH sensitivity.

In conclusion, we report a series of high-performance intracellular L-lactate biosensors that can be used to visualize intracellular L-lactate dynamics with large fluorescence responses and over a wide concentration range. We expect that the iLACCO series should be highly amenable to a broad range of further applications, including *in vivo* imaging to investigate physiological L-lactate concentration dynamics in animal models.

## Materials and Methods

### General methods and materials

A synthetic human codon-optimized gene encoding the *Escherichia coli* LldR transcriptional regulator protein was purchased from Integrated DNA Technologies. Phusion high-fidelity DNA polymerase (Thermo Fisher Scientific) was used for routine polymerase chain reaction (PCR) amplification, and Taq DNA polymerase (New England Biolabs) was used for error-prone PCR. QuickChange mutagenesis kit (Agilent Technologies) was used for site-directed mutagenesis. Restriction endonucleases and rapid DNA ligation kits (Thermo Fisher Scientific) were used for plasmid construction. Products of PCR and restriction digests were purified using agarose gel electrophoresis and the GeneJET gel extraction kit (Thermo Fisher Scientific). DNA sequences were analyzed by DNA sequence service of Fasmac Co. Ltd. Fluorescence spectra and intensity were recorded on Spark plate reader (Tecan) or CLARIOstar Plus Microplate reader (BMG LABTECH).

### Structural modeling of LldR and iLACCO1

The modeling structure of LldR and iLACCO1 was generated by AlphaFold2^31, 32^ using an API hosted at the Södinglab in which the MMseqs2 server^40^ was used for multiple sequence alignment. The amino acid sequence of *Escherichia coli* LldR transcriptional regulator protein was submitted to ColabFold to generate the modeling structure of LldR^32^. The amino acid sequence of iLACCO0.2 was submitted to ColabFold to generate the template for iLACCO1 structure. This specific sequence was chosen because it does not include newly introduced mutations which make the sequence identity closer to what can be found in the training data set of Alphafold2.

### Engineering of iLACCO1 variants

The gene encoding cpGFP with N- and C-terminal linkers (DWS and NDG, respectively) was amplified, followed by insertion into each site of LldR-LBD in a pBAD vector (Life Technologies) by Gibson assembly (New England Biolabs). DNA binding domain of LldR was removed beforehand. Variants were expressed in *E. coli* strain DH10B (Thermo Fisher Scientific) in LB media supplemented with 100 µg mL^-1^ ampicillin and 0.02% L-arabinose. Proteins were extracted by B-PER bacterial protein extraction reagent (Thermo Fisher Scientific) for the assay of fluorescence brightness and L-lactate-dependent response at screening. During evolution processes, MnCl_2_ concentration was controlled to obtain one to two mutations in the whole gene per round. Primary screening was done on the agar plates, where approximately 2 × 10^3^ colonies were visually inspected each round. Total 192 bright colonies were then picked up for protein extraction and fluorescence measurement. After three rounds of linker optimization, 11 rounds of directed evolution in the whole gene sequence followed by the introduction of C164S mutation ultimately led to iLACCO1. Mutations for tuning affinity were introduced by site-directed mutagenesis using the QuikChange mutagenesis kit.

### Protein purification

iLACCO variants in pBAD expression vectors containing a N-terminal poly His (6×) tag were expressed in *E. coli* BL21(DE3). A single colony was used to inoculate a 10 mL culture of 100 µg mL^-1^ ampicillin LB and grown overnight at 37 °C by shaking at 180 rpm. One mL of saturated culture was then used to inoculate 1 L of LB supplemented with 100 µg mL^-1^ ampicillin and grown at 37 °C until OD_600_ value of 0.6 was reached. The cell culture was then induced by adding 0.1% [w/v] L-arabinose and grown overnight at 18 °C. The next day, cells were harvested by centrifugation at 4 °C and 4,000 rpm for 1h. Each 1 g of cell pellet was resuspended in 6 mL of lysis buffer (30 mM MOPS, 100 mM KCl, 10% [v/v] glycerol, 1 mM TCEP, 1 mM PMSF, 5 mM benzamidine, 10 mM imidazole, pH 7.2). Cells were placed on ice and lysed by sonication (30 s on/off for 2.5 min, 2.5 min off, and 30 s on/off for 2.5 min) and then centrifuged at 12,000 rpm and 4 °C for 1h. Lysates were filtered with a 0.45 µm filter and loaded onto a lysis buffer pre-equilibrated Ni-NTA column. The column was then washed with 10 column volume (cv) of lysis buffer. Protein was eluted with 3 cv of elution buffer (30 mM MOPS, 100 mM KCl, 2% [v/v] glycerol, 1 mM TCEP, 0.2 mM PMSF, 0.2 mM benzamidine, 500 mM imidazole, pH 7.2) and buffer exchanged into L-lactate (-) buffer (30 mM MOPS, 100 mM KCl, pH 7.2) using centrifugal spin column (10K MWCO, Thermo Fisher Scientific). Prior to analysis, all proteins were then further purified by size exclusion chromatography using a Superdex 75 10/300 GL increase column (GE Healthcare).

### *In vitro* characterization

Absorption spectra were collected on SPECTROstar Nano microplate reader (BMG LABTECH) using 10 mm quartz cuvette (Hellma Analytics). To perform rapid kinetic measurement for the interaction between iLACCO1 and L-lactate, absorption at 493 nm was detected through a 10 mm quartz cuvette using a diode array UV-Vis spectrophotometer (Ocean Optic Inc., USB4000). All fluorescence spectra were collected on CLARIOstar Plus Microplate reader (BMG LABTECH) using Greiner 96-well Flat-bottom microplate. For absorption and fluorescence excitation/emission spectra, L-lactate (-) buffer and L-lactate (+) buffer (30 mM MOPS, 100 mM KCl, 100 mM L-lactate, pH 7.2) were used. To perform measurement of *K*_d_, a series of buffers with L-lactate concentration ranging from 0 to 100 mM was prepared by diluting L-lactate (+) buffer using L-lactate (-) buffer. Sensitivity of sensors as a function of L-lactate concentration was then fitted to the Hill equation (f(*x*) = base + (max - base)/(1 + (*K*_d_/*x*)^Hill^ ^coefficient^)) to determine the Hill coefficient and apparent *K*_d_. Sensitivity of sensors are reported as Δ*F*/*F* which is calculated by (*F*_x_ - *F*_(-)_)/*F*_(-)_, where *F*_x_ is the fluorescence intensity at 510 nm of sample x and *F*_(-)_ is the fluorescence intensity at 510 nm of same concentration of protein in L-lactate (-) buffer. For pH titration, pH buffer (30 mM MOPS, 30 mM trisodium citrate, 30 mM sodium borate, 100 mM KCl, and either no L-lactate or 10 mM L-lactate, pH ranging from 5 to 10) were used to dilute protein solutions. Fluorescence intensities as a function of pH were then fitted to a sigmoidal function (f(pH) = base + (max-base)/(1+10^p*K*a-pH^)) to determine the p*K*_a_. All measurements were conducted at room temperature.

### Two-photon absorption measurements

Two-photon excitation spectra and two-photon absorption cross sections were measured using standard methods and protocols^41^. Briefly, a tunable femtosecond laser InSight DeepSee (Spectra-Physics, Santa Clara, CA) was used to excite fluorescence of the sample in a PC1 Spectrofluorometer (ISS, Champaign, IL). To measure the two-photon excitation spectral shapes, we used in the emission channel a combination of short-pass filters 633SP and 770SP for iLACCO1 and iLACCO1.2, and an additional 535/50 filter for iLACCO1.1. Organic dyes LDS 798 in 1:2 CHCl_3_:CDCl_3_ and Coumarin 540A in DMSO were used as spectral standards. Quadratic power dependence of fluorescence intensity in the proteins and standards was checked at several wavelengths across the spectrum.

The two-photon cross section (σ_2_) of the anionic form of the chromophore was measured as described previously^42^. Fluorescein in water at pH 12 was used as a reference standard with excitation at 900 nm^41^. For one-photon excitation we used a 488-nm line of an argon ion laser (Melles Griot) and a combination of filters 770SP and 520LP was in the fluorescence channel. Extinction coefficients were determined by alkaline denaturation as previously described^43^. The two-photon absorption spectra were normalized to the measured σ_2_ values. To normalize to the total two-photon brightness (*F*_2_), the spectra were multiplied by the quantum yield and the relative fraction of the anionic form of the chromophore. The data is presented this way because iLACCO1, iLACCO1.1, and iLACCO1.2 all exist as mixtures of the neutral and anionic forms of the chromophore at neutral pH. The method has been previously described in detail^43^.

### Construction of mammalian expression vectors

The gene of an iLACCO variant was amplified with sequence coding P2A self-cleaving peptide by PCR and cut with XhoⅠ and EcoRⅠ. The gene encoding pHuji^36^ was amplified by PCR followed by digestion with EcoRI and HindIII. Finally, these products were ligated into pcDNA3 vector (Thermo Fisher Scientific) with T4 ligase (Thermo Fisher Scientific). The gene of Laconic (Addgene plasmid no. 44238) and Green Lindoblum (synthetic DNA purchased from Integrated DNA Technologies) were ligated into a pcDNA vector without pHuji.

### Imaging of iLACCO variants in HeLa and HEK293 cells

HeLa (Japanese Cancer Research Resources Bank; JCRB and American Type Culture Collection; ATCC) and HEK293 (JCRB) cells were maintained in Dulbecco’s modified Eagle medium (DMEM high glucose; Nacalai Tesque) supplemented with 10% fetal bovine serum (FBS; Sigma-Aldrich), and 100 μg mL^-1^ penicillin and streptomycin (Nacalai Tesque). Cells were transiently transfected with the plasmids with polyethyleneimine (Polysciences) in Opti-MEM (Gibco) and imaged within 48–72 hours after transfection. IX83 wide-field fluorescence microscopy (Olympus) equipped with a pE-300 LED light source (CoolLED), and a 40× objective lens (numerical aperture (NA) = 1.3; oil), an ImagEM X2 EM-CCD camera (Hamamatsu), and Cellsens software (Olympus) were used for the imaging. The filter sets in the imaging were the following specification. iLACCO1 variants and Green Lindoblum: excitation 470/20 nm, dichroic mirror 490 nm dclp, and emission 518/45 nm; pHuji: excitation 545/20 nm, dichroic mirror 565 nm dclp, and emission 598/55 nm; Laconic (mTFP1): excitation 438/24 nm, dichroic mirror 458 nm dclp, and emission 483/32 nm; Laconic (FRET): excitation 438/24 nm, dichroic mirror 458 nm dclp, and emission 542/27 nm. Fluorescence images were analyzed with ImageJ software (https://imagej.net/software/fiji/, National Institutes of Health).

In starvation experiments, HeLa and HEK293 cells were incubated in no-glucose DMEM (Nacalai Tesque) for 2–3 hours before imaging. After exchanging medium into no-glucose imaging buffer (184.45 mg L^-1^ CaCl_2_·H_2_O, 97.6 mg L^-1^ MgSO_4_, 400.00 mg L^-1^ KCl, 60.00 mg L^-1^ KH_2_PO_4_, 8000.00 mg L^-1^ NaCl, 350.00 mg L^-1^ NaHCO_3_, 47.88 mg L^-1^ Na_2_HPO_4_), final concentration of 5 mM D-glucose was added at time = 0 min.

For imaging in treatment with MCT inhibitor AR-C155858 (Tocris), Hank’s balanced salt solution (HBSS; Nacalai Tesque) supplemented with additional D-glucose (final concentration of 25 mM) and 10 mM HEPES (Nacalai Tesque) was used as an imaging buffer.

### Flow cytometry of HeLa cells expressing iLACCO variants

Within 48–72 hours after the transfection, cells were collected after incubation with 500 µM iodoacetic acid (the same amount of MilliQ or none was added into control samples), and washed with PBS. The cells were suspended into HBSS supplemented with 10 mM HEPES and respective reagents (10 µM nigericin and 2 µM rotenone into stimulated samples) and were passed through a cell-strainer with 35 μm-mesh (Falcon). Flow cytometry analysis was carried out using SH800 (Sony). The data were analyzed by FlowJo software (BD).

## Data availability

The data and plasmids encoding iLACCO variants that support the findings of this study are available from the corresponding authors on reasonable request.

## Acknowledgments

The authors thank T. Terai for technical support. We thank K. Miyazono, G.A. Woolley and R. Tanabe for providing access to their instruments and technical support. Work at the University of Tokyo was supported by the Japan Society for the Promotion of Science (Y.N., Grants-in-Aid for Early-Career Scientists 21K14738; R.E.C., Grants-in-Aid for Scientific Research S 19H05633). G.N.T.L. is supported by the Global Science Course (GSC) program and NSERC CREATE Advanced Protein Engineering Training, Internships, Courses, and Exhibition (APRENTICE) program.

## Author Contributions

G.N.T.L. and S.L. developed iLACCO1. S.H. developed iLACCO affinity variants. G.N.T.L. and S.H. performed *in vitro* characterization. M.D. measured two-photon excitation spectra. S.H. and K.T-Y. performed FACS experiment, live cell imaging, and data analysis. Y.N. and R.E.C. supervised research. G.N.T.L., S.H., K.T-Y., M.D., Y.N., and R.E.C. wrote the manuscript.

## Supplementary Information

### Supplementary Figures

**Supplementary Figure 1.**
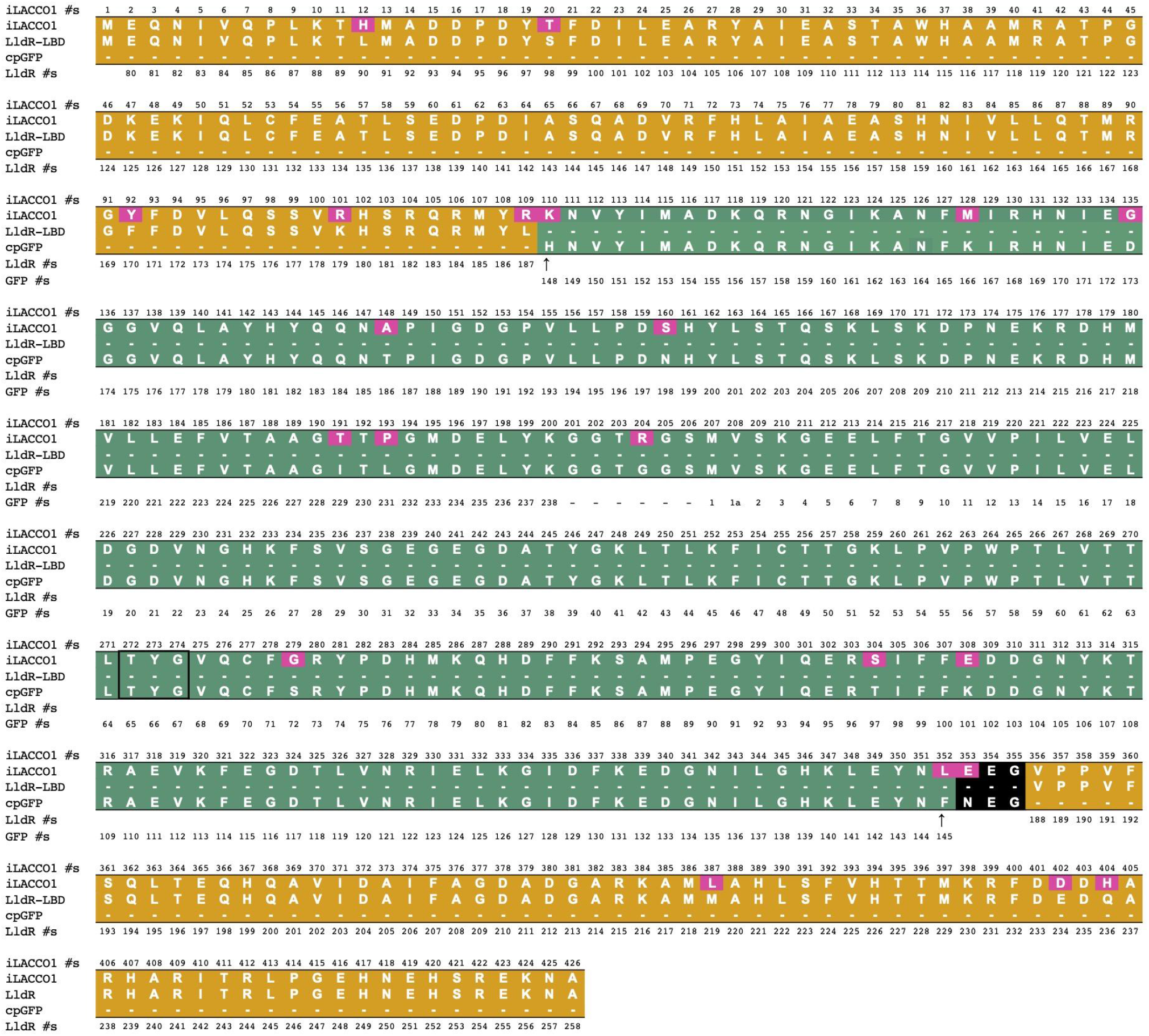
Sequence alignment of iLACCO1. The LldR-LBD domain is indicated by an orange background and the cpGFP domain is indicated by a green background. Mutations in iLACCO1, relative to LldR-LBD and cpGFP, are indicated by a magenta background. The linker at the C-terminal end of cpGFP is indicated by a black background. The gate post residues at the ends of cpGFP (residues 145 and 148), are indicated by upward pointing arrows. The linker joining the original C- and N-termini of GFP (residues 238 and 1, respectively) is not numbered. The chromophore-forming residues are enclosed in black box.

**Supplementary Figure 2.**
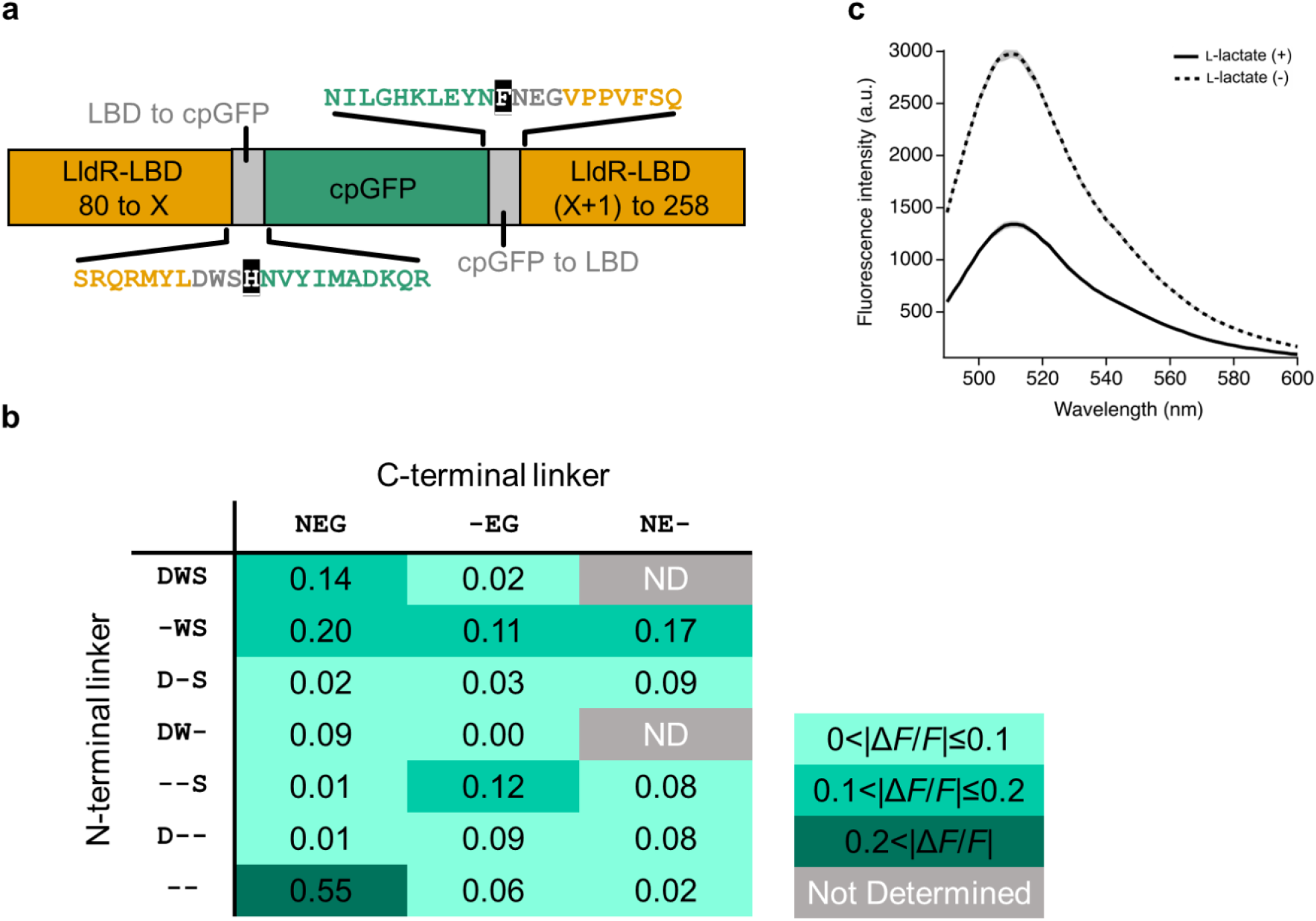
Optimization of linker lengths. **a)** Schematic representation of the primary structure of iLACCO0.1. iLACCO0.1 has DWS and NEG as N- and C-terminal linker, respectively. **b)** Δ*F*/*F* profile of each linker length variant. The variant with no N-terminal linker and original C-terminal linker residues (NEG) showed the largest absolute value of Δ*F*/*F* and the protein was designated iLACCO0.2. **c)** Emission spectra of iLACCO0.2 in the presence (10 mM, represented in solid line) and absence of L-lactate (represented in dashed line).

**Supplementary Figure 3.**
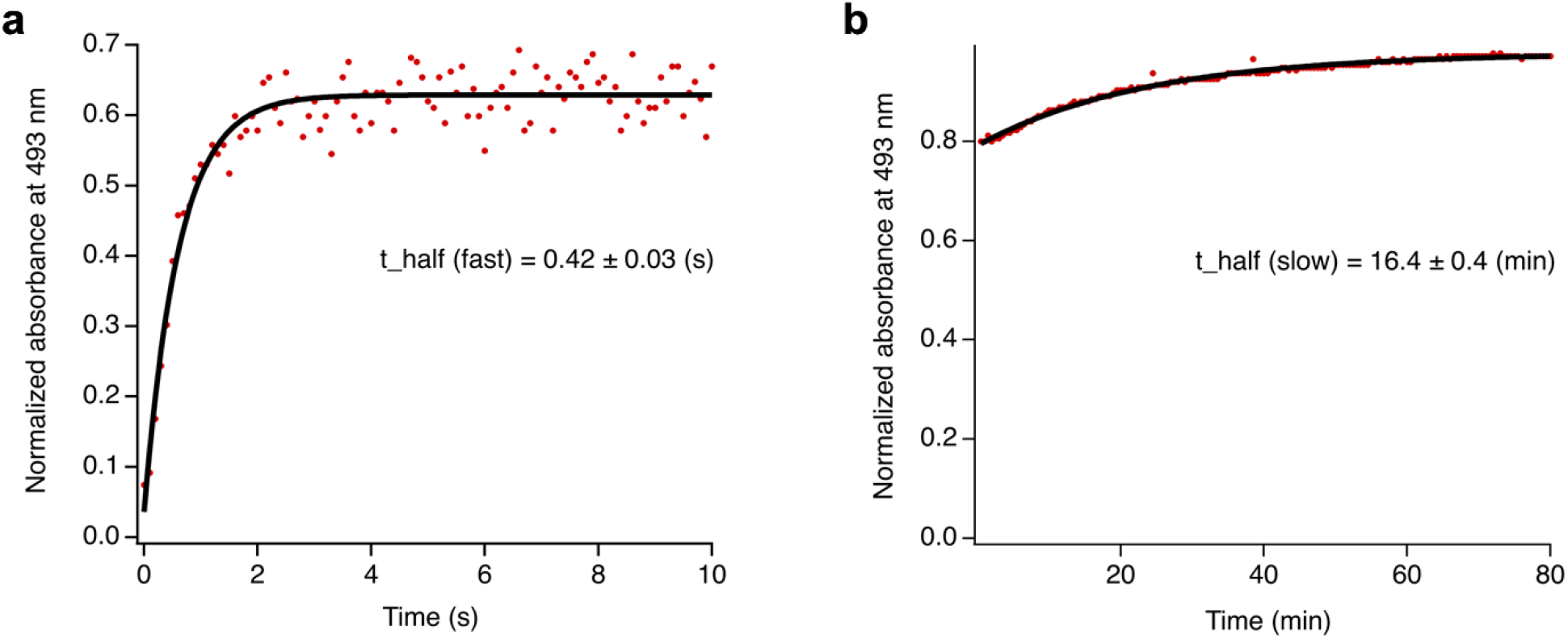
Kinetics of iLACCO1 *in vitro*. Normalized absorbance at 493 nm or purified iLACCO1 protein after adding 10 mM L-lactate as a function of time. The data was fitted by f(t) = c + A * (1-exp(-(ln(2)/t_half)*t)). **a)** Absorption data was collected every 100 ms for 10 s, integration time: 30 ms, scans to average: 3, boxcar width: 25. **b)** Absorption data was collected every 30 s for 80 minutes.

**Supplementary Figure 4.**
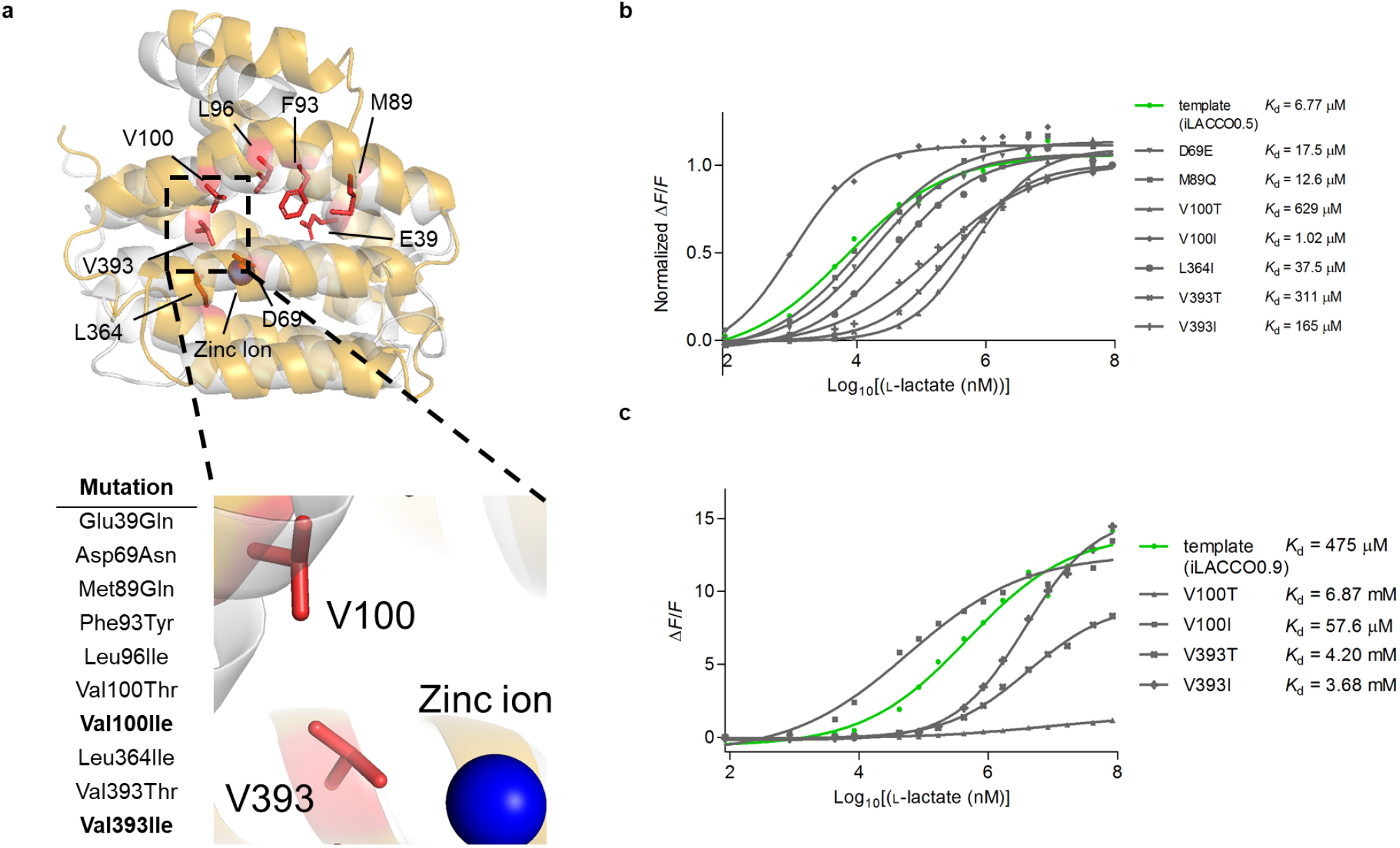
Engineering of affinity variants of iLACCO. **a)** A homology model of LldR from *E. coli* (based on LldR from *Corynebacterium glutamicum*; PDB ID: 2DI3)^1^ calculated on M4T Server ver 3.0 (ref. 2) (yellow), and superpositioned with the crystal structure of LldR from *C. glutamicum* (gray). Amino acid residues in the binding pocket that were mutated for possible affinity tuning are shown in red. The blue sphere indicates a Zn^2+^. **b)** Dose-response curves and calculated *K*_d_ values for mutants of iLACCO0.5 (*n* = 1, the same trend was confirmed in independent experiments). Three mutations (V100T, V393T, and V393I) that decreased the affinity and one mutation (V100I) that increased the affinity were selected for testing on later variants in directed evolution. **c)** Dose-response curves and calculated *K*_d_ and Δ*F*/*F* values for mutated variants of iLACCO0.9 (n = 1, the same trend was confirmed in independent experiments). V100I was applied for tuning affinity higher and V393I was applied for tuning affinity lower. These mutations were later introduced to the final variant iLACCO1.

**Supplementary Figure 5.**
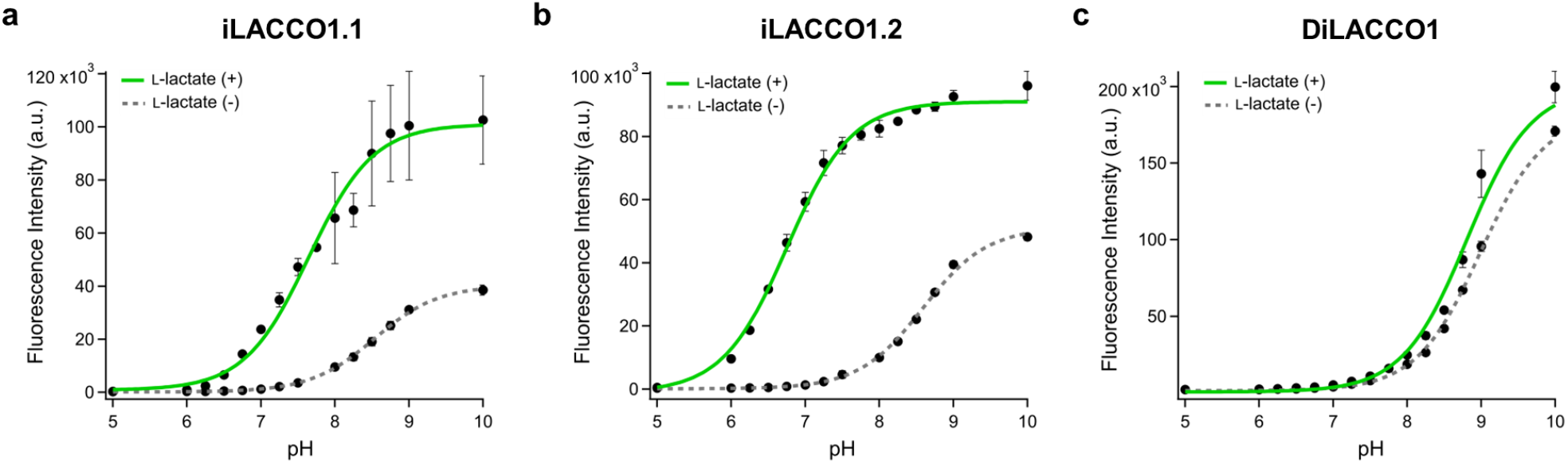
pH titration curve of iLACCO variants. pH titration curve of **a)** iLACCO 1.1, **b)** iLACCO1.2, and **c)** DiLACCO1 in the presence (10 mM) and absence of L-lactate. *n* = 3 technical replicates (mean ± s.d.).

**Supplementary Figure 6.**
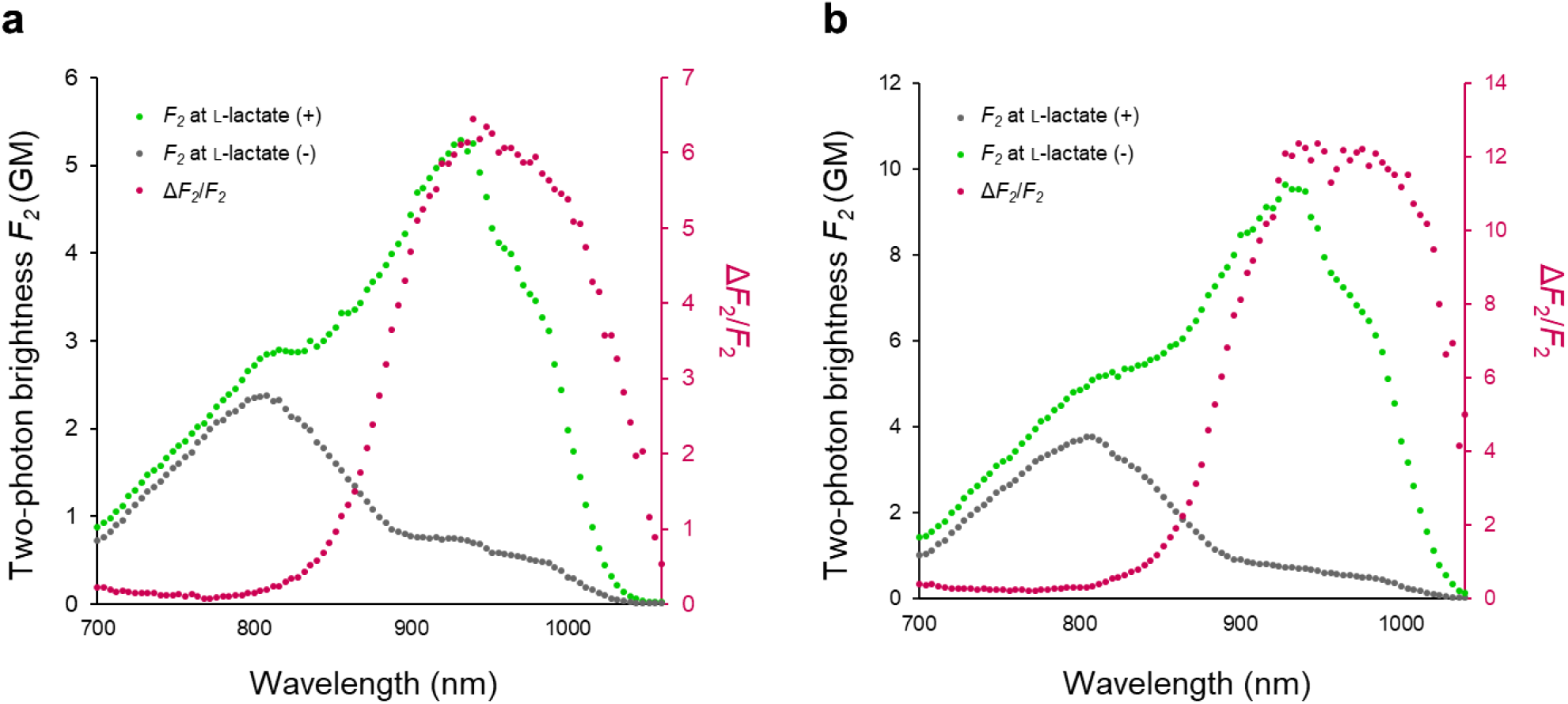
Two-photon spectra of iLACCO1.1 and iLACCO1.2. Two-photon spectra of a) iLACCO1.1 and b) iLACCO1.2.

**Supplementary Figure 7.**
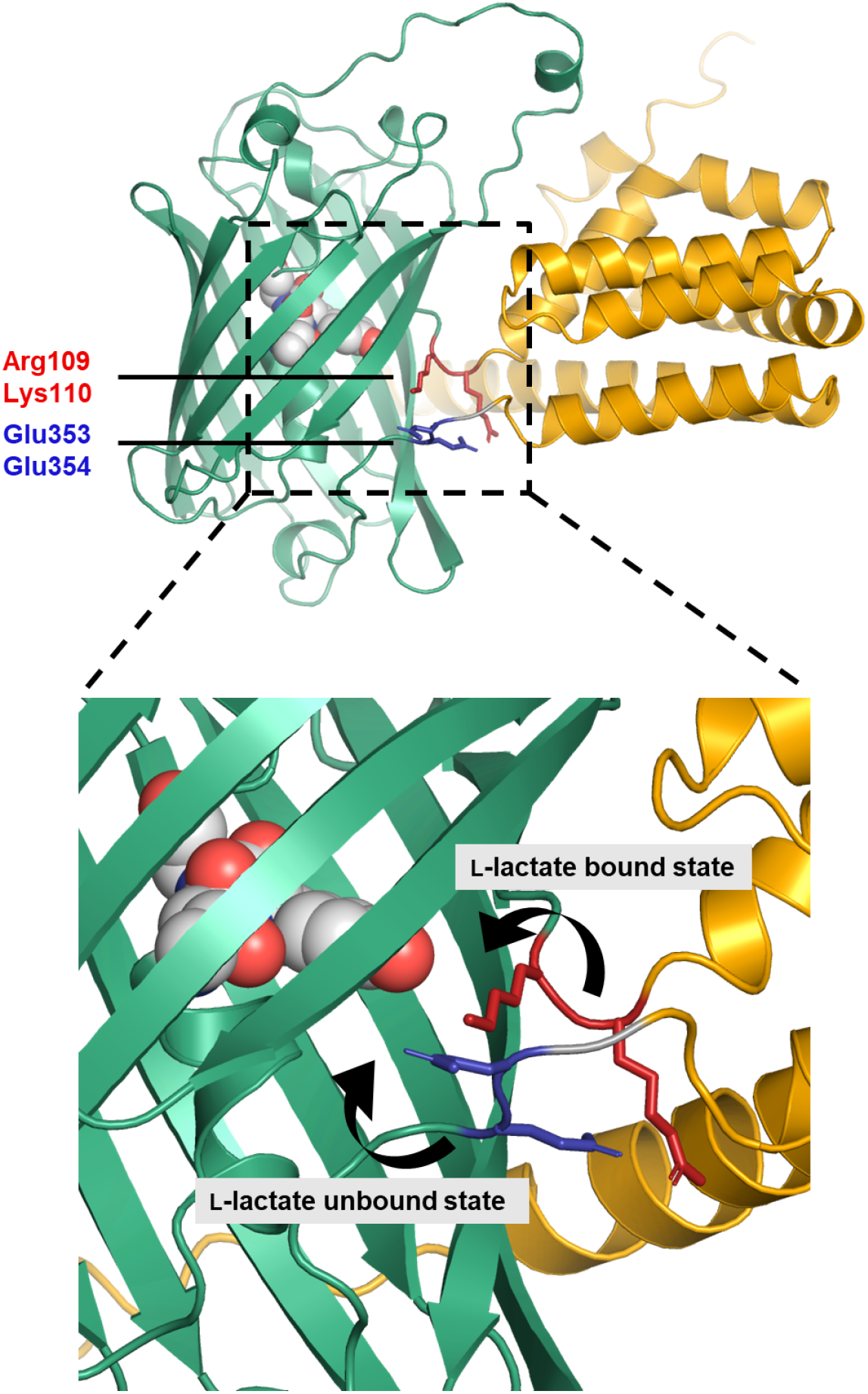
Proposed response mechanism and chromophore interactions of iLACCO1. Shown is an Alphafold model of iLACCO1 with the chromophore-forming tripeptide represented as spheres, the N-terminal gate post and its preceding residue (His110Lys and Leu109Arg, respectively) represented as red sticks, and the C-terminal gate post and its following residue (Asn353Glu and Glu354, respectively) represented as blue sticks. As described in the main text, we tentatively suggest that in the L-lactate-bound state, the positively charged residues shown in red are interacting with the chromophore and stabilizing in the anionic phenolate state. Conversely, in the L-lactate unbound state, the negatively charged residues shown in blue may be interacting with the chromophore and stabilizing it in the neutral phenol state.

### Supplementary Tables

**Supplementary Table 1.** Photophysical and biochemical properties of the iLACCO series.

| | Key Mutations | $\Delta F/F$ | $K_d$ (mM) | +/- Lac | Excitation peak (nm) | Emission peak (nm) | p*K*<sub>a</sub> | Anionic QY |
| --- | --- | --- | --- | --- | --- | --- | --- | --- |
| iLACCO1 | NA | 29.7 ± 0.5 | 0.36 ± 0.03 | + | 493 | 510 | 7.38 ± 0.06 | 0.84 ± 0.01 |
|  |  |  |  | - | 492 | 511 | 8.78 ± 0.03 | 0.72 ± 0.03 |
| iLACCO1.1 | V393I | 15.2 ± 0.4 | 4.6 ± 1.0 | + | 493 | 510 | 7.65 ± 0.06 | 0.57 ± 0.07 |
|  |  |  |  | - | 493 | 510 | 8.53 ± 0.01 | 0.57 ± 0.07 |
| iLACCO1.2 | V100I | 28.4 ± 2.7 | 0.017 ± 0.002 | + | 493 | 509 | 6.76 ± 0.03 | 0.86 ± 0.01 |
|  |  |  |  | - | 493 | 510 | 8.59 ± 0.02 | 0.61 ± 0.03 |
| DiLACCO1 | V100A<br>V393A | 0.4 ± 0.2 | NA | + | 493 | 510 | 8.81 ± 0.05 | ND |
|  |  |  |  | - | 493 | 510 | 8.99 ± 0.02 | ND |
| QY, quantum yield.<br>NA, not applicable.<br>ND, not determined.<br>*n* = 3 technical replicates (mean ± s.d.). |  |  |  |  |  |  |  |  |

## References

1. Levitt, M. D. & Levitt, D. G. Quantitative Evaluation of D-Lactate Pathophysiology: New Insights into the Mechanisms Involved and the Many Areas in Need of Further Investigation. Clin. Exp. Gastroenterol. 13, 321–337 (2020).

2. Hill, A. V. & Kupalov, P. Anaerobic and aerobic activity in isolated muscle. Proc. R. Soc. Lond. B Biol. Sci. 105, 313–322 (1929).

3. Rabinowitz, J. D. & Enerbäck, S. Lactate: the ugly duckling of energy metabolism. Nat Metab 2, 566–571 (2020).

4. Brooks, G. A. Lactate as a fulcrum of metabolism. Redox Biol 35, 101454 (2020).

5. Magistretti, P. J. & Allaman, I. Lactate in the brain: from metabolic end-product to signalling molecule. Nat. Rev. Neurosci. 19, 235–249 (2018).

6. Mosienko, V., Teschemacher, A. G. & Kasparov, S. Is L-lactate a novel signaling molecule in the brain? J. Cereb. Blood Flow Metab. 35, 1069–1075 (2015).

7. Suzuki, A. et al. Astrocyte-neuron lactate transport is required for long-term memory formation. Cell 144, 810–823 (2011).

8. Ratter, J. M. et al. In vitro and in vivo Effects of Lactate on Metabolism and Cytokine Production of Human Primary PBMCs and Monocytes. Front. Immunol. 9, 2564 (2018).

9. Angelin, A. et al. Foxp3 Reprograms T Cell Metabolism to Function in Low-Glucose, High-Lactate Environments. Cell Metab. 25, 1282–1293.e7 (2017).

10. Lev-Vachnish, Y. et al. L-Lactate Promotes Adult Hippocampal Neurogenesis. Front. Neurosci. 13, 403 (2019).

11. Pucino, V., Bombardieri, M., Pitzalis, C. & Mauro, C. Lactate at the crossroads of metabolism, inflammation, and autoimmunity. Eur. J. Immunol. 47, 14–21 (2017).

12. Haas, R. et al. Lactate Regulates Metabolic and Pro-inflammatory Circuits in Control of T Cell Migration and Effector Functions. PLoS Biol. 13, e1002202 (2015).

13. Warburg, O., Wind, F. & Negelein, E. The metabolism of tumors in the body. J. Gen. Physiol. 8, 519–530 (1927).

14. de la Cruz-López, K. G., Castro-Muñoz, L. J., Reyes-Hernández, D. O., García-Carrancá, A. & Manzo-Merino, J. Lactate in the Regulation of Tumor Microenvironment and Therapeutic Approaches. Front. Oncol. 9, 1143 (2019).

15. Hirschhaeuser, F., Sattler, U. G. A. & Mueller-Klieser, W. Lactate: a metabolic key player in cancer. Cancer Res. 71, 6921–6925 (2011).

16. Watson, M. J. et al. Metabolic support of tumour-infiltrating regulatory T cells by lactic acid. Nature 591, 645–651 (2021).

17. Frame, A. K., Simon, A. F. & Cumming, R. C. Determining the role of lactate metabolism on age-dependent memory decline and neurodegeneration in Drosophila melanogaster. Alzheimers. Dement. 16, (2020).

18. San Martín, A., et al. A genetically encoded FRET lactate sensor and its use to detect the Warburg effect in single cancer cells. PLoS One 8, e57712 (2013).

19. Warburg, O. The metabolism of carcinoma cells. J. Cancer Res. 9, 148–163 (1925).

20. Mächler, P. et al. In Vivo Evidence for a Lactate Gradient from Astrocytes to Neurons. Cell Metab. 23, 94–102 (2016).

21. Pellerin, L. & Magistretti, P. J. Glutamate uptake into astrocytes stimulates aerobic glycolysis: a mechanism coupling neuronal activity to glucose utilization. Proc. Natl. Acad. Sci. U. S. A. 91, 10625–10629 (1994).

22. Harada, K. et al. Green fluorescent protein-based lactate and pyruvate indicators suitable for biochemical assays and live cell imaging. Sci. Rep. 10, 19562 (2020).

23. Bekdash, R. et al. GEM-IL: A highly responsive fluorescent lactate indicator. Cell Rep Methods 1, 100092 (2021).

24. Galaz, A. et al. Highly responsive single-fluorophore indicator to explore lactate dynamics in high calcium environments. bioRxiv 2020.10.01.322404 (2020) doi:10.1101/2020.10.01.322404.

25. Koveal, D. et al. A high-throughput multiparameter screen for accelerated development and optimization of soluble genetically encoded fluorescent biosensors. Nat. Commun. 13, 2919 (2022).

26. Nasu, Y. et al. A genetically encoded fluorescent biosensor for extracellular L-lactate. Nat. Commun. 12, 7058 (2021).

27. Nasu, Y. et al. A red fluorescent genetically encoded biosensor for extracellular L-lactate. bioRxiv 2022.08.30.505811 (2022) doi:10.1101/2022.08.30.505811.

28. Nasu, Y., Shen, Y., Kramer, L. & Campbell, R. E. Structure- and mechanism-guided design of single fluorescent protein-based biosensors. Nat. Chem. Biol. 17, 509–518 (2021).

29. Marvin, J. S. et al. An optimized fluorescent probe for visualizing glutamate neurotransmission. Nat. Methods 10, 162–170 (2013).

30. Gao, Y.-G. et al. Structural and functional characterization of the LldR from Corynebacterium glutamicum: a transcriptional repressor involved in L-lactate and sugar utilization. Nucleic Acids Res. 36, 7110–7123 (2008).

31. Jumper, J. et al. Highly accurate protein structure prediction with AlphaFold. Nature 596, 583–589 (2021).

32. Mirdita, M. et al. ColabFold: making protein folding accessible to all. Nat. Methods 19, 679–682 (2022).

33. Brandt, R. B., Siegel, S. A., Waters, M. G. & Bloch, M. H. Spectrophotometric assay for D-(-)-lactate in plasma. Anal. Biochem. 102, 39–46 (1980).

34. Ewaschuk, J. B., Naylor, J. M. & Zello, G. A. D-lactate in human and ruminant metabolism. J. Nutr. 135, 1619–1625 (2005).

35. Fernandez-Fuentes, N., Rai, B. K., Madrid-Aliste, C. J., Fajardo, J. E. & Fiser, A. Comparative protein structure modeling by combining multiple templates and optimizing sequence-to-structure alignments. Bioinformatics 23, 2558–2565 (2007).

36. Shen, Y., Rosendale, M., Campbell, R. E. & Perrais, D. pHuji, a pH-sensitive red fluorescent protein for imaging of exo- and endocytosis. J. Cell Biol. 207, 419–432 (2014).

37. Szymczak-Workman, A. L., Vignali, K. M. & Vignali, D. A. A. Design and construction of 2A peptide-linked multicistronic vectors. Cold Spring Harb. Protoc. 2012, 199–204 (2012).

38. Ovens, M. J., Davies, A. J., Wilson, M. C., Murray, C. M. & Halestrap, A. P. AR-C155858 is a potent inhibitor of monocarboxylate transporters MCT1 and MCT2 that binds to an intracellular site involving transmembrane helices 7–10. Biochem. J 425, 523–530 (2010).

39. Poole, R. C. & Halestrap, A. P. Transport of lactate and other monocarboxylates across mammalian plasma membranes. Am. J. Physiol. 264, C761–82 (1993).

40. Steinegger, M. & Söding, J. MMseqs2 enables sensitive protein sequence searching for the analysis of massive data sets. Nat. Biotechnol. 35, 1026–1028 (2017).

41. Drobizhev, M., Molina, R. S. & Hughes, T. E. Characterizing the Two-photon Absorption Properties of Fluorescent Molecules in the 680-1300 nm Spectral Range. Bio Protoc 10, (2020).

42. Dalangin, R. et al. Far-red fluorescent genetically encoded calcium ion indicators. bioRxiv 2020.11.12.380089 (2020) doi:10.1101/2020.11.12.380089.

43. Barnett, L. M., Hughes, T. E. & Drobizhev, M. Deciphering the molecular mechanism responsible for GCaMP6m’s Ca2+-dependent change in fluorescence. PLoS One 12, e0170934 (2017).

44. Kalivoda, K. A., Steenbergen, S. M. & Vimr, E. R. Control of the Escherichia coli sialoregulon by transcriptional repressor NanR. J. Bacteriol. 195, 4689–4701 (2013).

## Supplementary References

1. Gao, Y.-G. et al. Structural and functional characterization of the LldR from Corynebacterium glutamicum: a transcriptional repressor involved in L-lactate and sugar utilization. Nucleic Acids Res. 36, 7110–7123 (2008).

2. Fernandez-Fuentes, N., Rai, B. K., Madrid-Aliste, C. J., Fajardo, J. E. & Fiser, A. Comparative protein structure modeling by combining multiple templates and optimizing sequence-to-structure alignments. Bioinformatics 23, 2558–2565 (2007).

